# Human adenylyl cyclase 9 is auto-stimulated by its isoform-specific C-terminal domain and equipped with paradigm-switching mechanisms

**DOI:** 10.1101/2022.06.05.494885

**Authors:** Zhihao Chen, Ferenc A. Antoni

**Author notes:** **Corresponding author:** Ferenc A. Antoni, Centre for Discovery Brain Sciences, Deanery of Biomedical Sciences, University of Edinburgh, 15 George Sq, Edinburgh EH8 9XD, Scotland, U.K., mailto.

## Abstract

Human transmembrane adenylyl cyclase 9 (AC9) is not regulated by heterotrimeric G proteins. Key to the resistance to stimulation by Gs-coupled receptors (GsR) is auto-inhibition by the COOH-terminal domain (C2b). The present study investigated the role of the C2b domain in the regulation of cyclic AMP production by AC9 in HEK293FT cells expressing the Glosensor22F cyclic AMP-reporter protein. Surprisingly, we found C2b to be essential for sustaining the basal output of cyclic AMP by AC9. A human mutation (E326D) in the parallel coiled-coil formed by the signalling helices of AC9 dramatically increased basal activity, which was also dependent on the C2b domain. Intriguingly, the same mutation enabled stimulation of AC9 by GsRs. In summary, auto-regulation by the C2b domain of AC9 sustains its basal activity and quenches activation by GsR. Thus AC9 appears to be tailored to support constitutive activation of cyclic AMP effector systems. A switch from this paradigm to stimulation by GsRs may be occasioned by conformation changes at the coiled-coil or removal of the C2b domain.

## Introduction

Transmembrane adenylyl cyclases produce the ubiquitous signalling molecule adenosine-3’:5’-monophosphate (cAMP). Nine genes encode these enzymes in mammals and each paralogue has unique regulatory properties (Ostrom *et al*, 2022). Adenylyl cyclase 9 (AC9) is widely distributed in the body and has been implicated in a number of physiologic processes, including cardiac function, body fat mass and body weight, as well as cancer pathologies and atherosclerosis - (Antoni, 2020; Ostrom *et al*., 2022). Partial high-resolution maps of the structure of AC9 obtained by cryo-electronmicroscopy (cryoEM) have been recently published (Qi *et al*, 2022; Qi *et al*, 2019). In brief, the 1353-residue single polypeptide chain of AC9 forms a tripartite structure. This consists of a large trans-membrane array that is connected to the catalytic domain in the cytoplasm by two alpha helices (Fig 1A).

**Figure 1.**
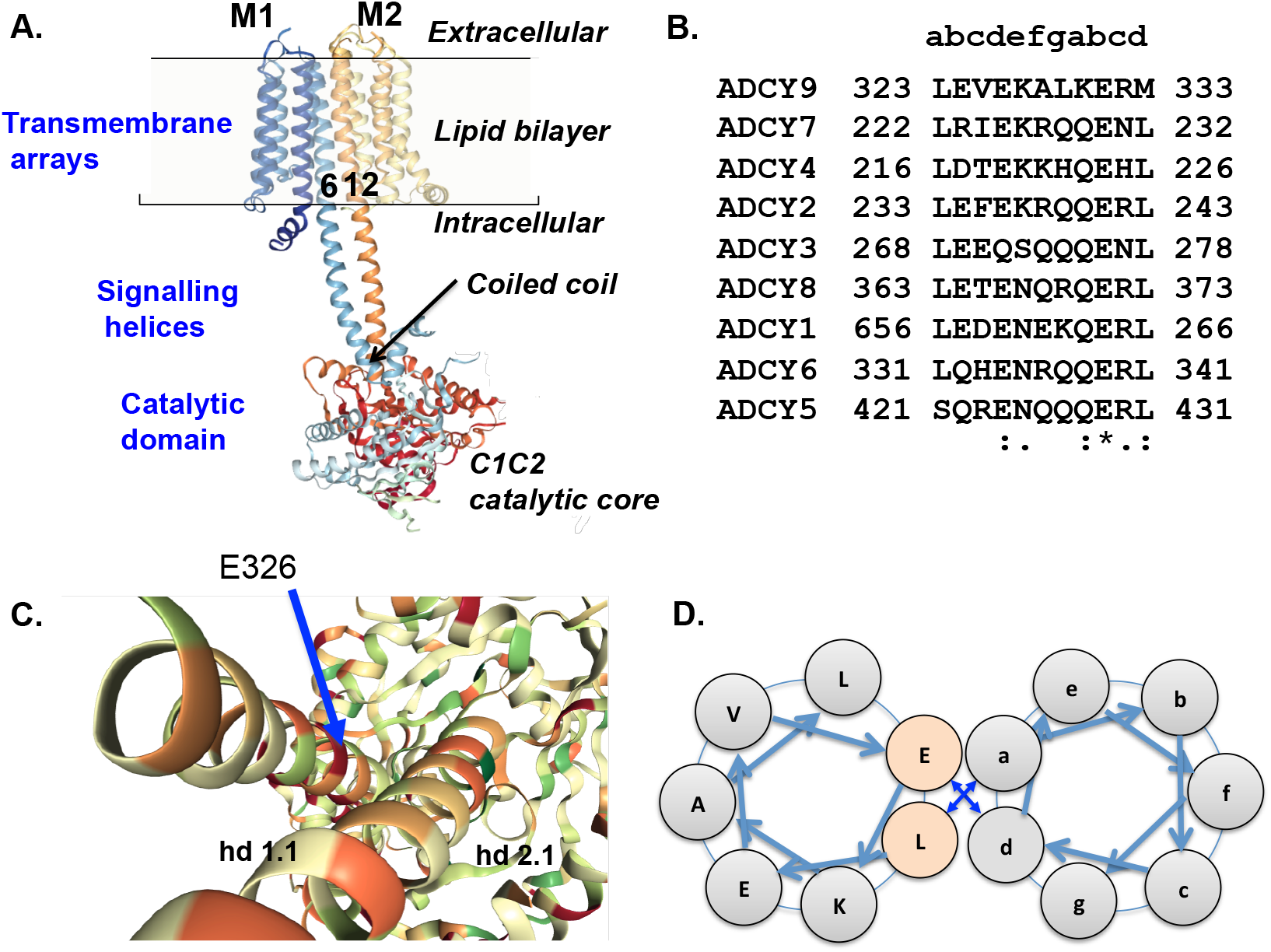
A) The tripartite structure of AC9 derived from cryo-EM studies by Qi and co-workers (Qi *et al*., 2022; Qi *et al*., 2019). Note that the enzyme consists of a 1353-residue single polypeptide and that some parts of the protein (shown by dashed red line) have not been resolved in the cryo-EM studies (Antoni, 2020). Transmembrane helices 6 and 12 are interfaced in the lipid bilayer and continue into the cytoplasm as the signaling helices that form a short, parallel coiled-coil just above the catalytic domain; B) Alignment of the hd1.1 (Qi *et al*., 2022; Qi *et al*., 2019) segment of human adenylyl cyclase paralogues. The small lettering in the top row indicates the positions in the coiled-coiled heptad of bovine AC9; C) E326 is at the interface of the coiled-coil, at position *d* of the heptad repeat as schematically shown in D)

The helices form a short parallel coiled-coil in close proximity to the catalytic core (Fig 1A). Sequence alignments of human adenylyl cyclases indicate that the coiled-coil of the “signalling helices” (Anantharaman *et al*, 2006; Bassler *et al*, 2018) is a generic feature of these proteins (Fig 1B). Coiled-coils are important regulatory modules in several proteins (Lupas & Bassler, 2017) including soluble guanylyl cyclases where the binding of nitric oxide is transmitted to the catalytic domains through conformational changes of the signalling helix (Horst *et al*, 2019).

Significantly, the enzymatic activity of full-length AC9 is largely insensitive to heterotrimec G proteins (Baldwin *et al*, 2019; Pálvölgyi *et al*, 2018; Qi *et al*., 2019). In the case of Gi/o this is due to the lack of a suitable binding pocket (Baldwin *et al*., 2019). With respect to Gs, the isoform-specific carboxyl-terminal (C2b) domain exerts a seemingly paradoxical auto-inhibitory effect by occluding the active site in the presence of Gsα-GTP (Qi *et al*., 2019). Given the unique regulatory features of AC9, this study investigated further the role of the C2b domain and its potential interactions with the coiled-coil.

The effects of a previously reported missense mutation, E326D (Calebiro *et al*, 2016), on cAMP production by human AC9 were analysed. This mutation is at the interface of the coiled-coil (Fig 1C and D). The results showed the E326D mutation markedly increased basal cAMP production by AC9, which was largely dependent on the presence of the isoform-specific C2b domain. In parallel, the mutation reduced the efficacy of the C2b domain to quench the activation of AC9 by Gs-coupled receptors (GsR). Finally, we show that the basal activity of wild-type AC9 also requires the C2b domain.

## Results and Discussion

### Effect of the coiled-coil mutation

Mutation E326D caused a 10-fold increase of basal cAMP production as well as an enhancement of the cAMP response to isoproterenol (see Fig. 2A for exemplar traces, and statistical summary in Fig. 3). Consistent with previous results with a different type of assay (Pálvölgyi *et al*., 2018), no isoproterenol-induced cAMP response attributable to wild-type AC9 could be reliably discerned. Others reported that a mutation in the predicted coil-coil of AC5 (M1029K) led to an enhanced GsR-induced cAMP response (Doyle *et al*, 2019; Lupas & Bassler, 2017; Qi *et al*., 2019), however, no changes in basal cAMP levels were found. In the case of AC9, aspartate instead of glutamate at the coiled-coil interface (E326D) in all probability changes the conformation of the coil (Jonson & Petersen, 2001; Straussman *et al*, 2007). The mutation led to a large, close to ten-fold increase in basal cAMP levels. In parallel, the cAMP response to GsR activation appeared, indicating a release from the potent auto-inhibitory effect exerted by the C2b domain (Pálvölgyi *et al*., 2018; Qi *et al*., 2019). The presence of such an adenylyl cyclase in thyroid epithelial cells (Calebiro *et al*., 2016) is likely to lead to hypertrophy and hyperplasia. In the context of a second oncogenic mutation, it can support adenomatous hyperproliferation (Calebiro *et al*., 2016) or epithelial mesenchymal transition producing malignant tumour growth (Tan *et al*, 2018).

**Figure 2.**
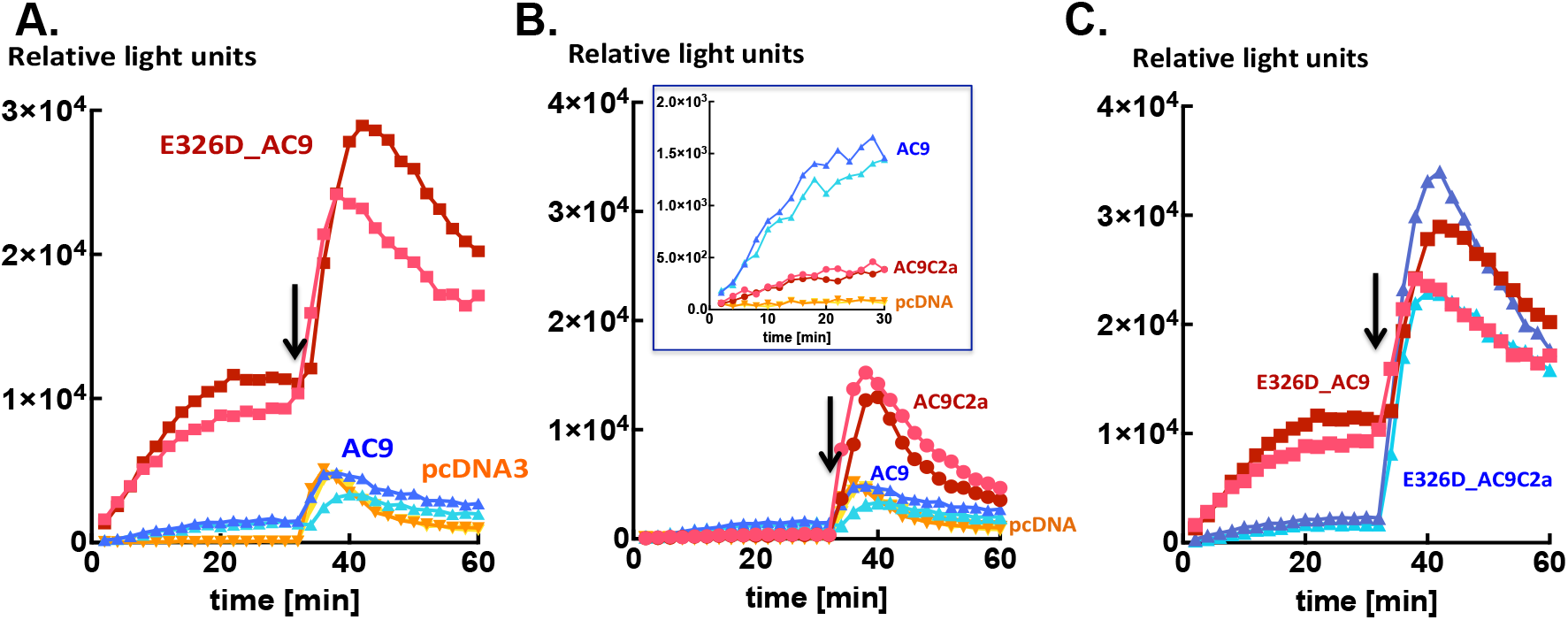
A) The mutation E326D markedly enhances cellular levels of cAMP. Intracellular levels of cAMP reported by Glosensor22F firefly luciferase bioluminescence in HEK293FT cells transiently transfected with AC9 (triangles (▴) blue hues), E326D_AC9 (square symbols (▪) red hues) or the skeleton vector pcDNA3 (wedges (▾) yellow hues). Isoproterenol (10 nM) was applied as indicated by the arrow. Single data-points recorded from individual wells of a 96-well microtiter plate are shown. The full data complement is shown in Fig 3. B) Effects of removing the isoform-specific C2b domain on intracellular levels of cAMP reported by Glosensor22F firefly luciferase bioluminescence from (A) HEK293FT cells transiently transfected with AC9 (triangles (▴) blue hues), AC9C2a (circles (•) red hues) or pcDNA3 (wedges (▾) yellow hues). The insert shows the basal cAMP levels from the same wells. The traces of AC9 and pcDNA3 are the ones already shown in Fig 2A; C) HEK293FT cells transiently transfected with E326D_AC9 (square symbols (▪) red hues) or E326D_AC9C2a (triangles (▴) blue hues). The traces of E326D_AC9 are the ones already shown in Fig. 2A. Isoproterenol (10 nM) was applied as indicated by the arrow. Single data points from individual wells of a 96-well microtiter plate. Data information: Representatives of six experiments from at least three different batches of transfected cells yielding closely similar results. The full data-set is shown in Fig. 3 and further similar experiments are presented in the Appendix.

**Figure 3.**
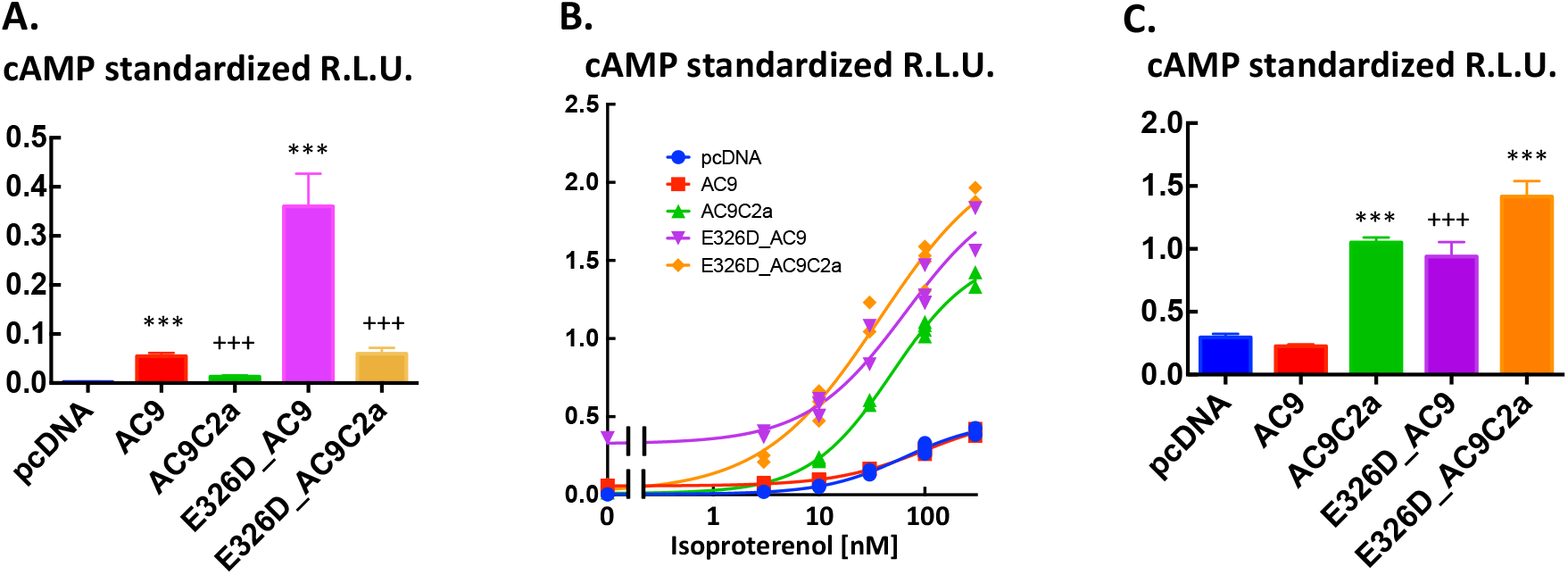
Auto-stimulation of basal and auto-inhibition of isoproterenol-evoked cAMP production by the C2b domain of AC9. Full data set of the exemplar traces shown in Fig 2. A) Basal intracellular cAMP levels reported by glosensorF22 bioluminescence in HEK293FT cells transiently expressing pcDNA3, AC9, AC9C2a, E326D_AC9 or, E326D_AC9C2a. Data are the average of the apparent plateau reached at 25-30 min of incubation (see Fig 2 for time courses) standardized in each well by the peak of the response elicited by 5μM forskolin and 100μM rolipram administered at the end of the experiment. Data are means ± S.D. n=14/group. As the variances of the five groups were statistically different, the data analysis was carried out after logtransformation of the data. One-way ANOVA F(4,65)=1464 P<0.0001, Tukey’s *post-hoc* multiple comparison test: *** P<0.001 *vs*. respective variant lacking the C2b domain. +++ P<0.0001 *vs*. pcDNA3 group. B) Concentration response to isoproterenol from the cells described in A). All measurements of the standardized peak responses are shown. Curves were fitted by non-linear four-parameter regression with variable slope in Graphpad Prism v.6. There was no difference between the EC50 values at alpha=0.05, the range was 37-63 nM. The maximal responses were E326D_AC9C2a= E326D_AC9=AC9C2a>AC9=pcDNA3 where > denotes statistical significance at P<0.05, as indicated by the 95% confidence intervals calculated by the non-linear regression algorithm. C) The increment over the respective basal levels induced by 100 nM isoproterenol calculated by averaging three consecutive time-points once the peak level of bioluminescence was reached (see Fig 2 for time-courses). The relative light-units were standardized in each well by the peak of the response elicited by 5μM forskolin and 100μM rolipram administered at the end of the experiment. Means ± S.D. N=4/group. One-way ANOVA F(4,15)=169.9 P<0.0001, Tukey’s *post-hoc* multiple comparison test: *** P<0.0001 *vs*. respective full-length variant. +++ P<0.0001 *vs*. pcDNA3 group. The results for 10 nM isoproterenol were closely similar.

### Probing the role of the C2b domain

Given the prominent role of the C2b domain in the regulation of the activity of AC9 (Pálvölgyi *et al*., 2018; Qi *et al*., 2019) we examined the effects of its deletion on cAMP levels produced by AC9 and E326D_AC9. Surprisingly, removal of the C2b domain from E326D_AC9 as well as wild-type AC9 reduced basal cAMP levels by 80-90% (see exemplar traces in Fig 2B and C, as well as and Fig 3 for the full dataset). With respect to stimulation by GsR, the amplitude of the agonist-induced cAMP response of AC9C2a was dramatically enhanced when compared with full-length AC9 (Fig 2B). This is fully consonant with previous results obtained by different methods of analysis (Pálvölgyi *et al*., 2018; Qi *et al*., 2019). However, it was not the case for E326D_AC9C2a: the peak levels of cAMP and the time-course of the isoproterenol response were not consistently different from those of full-length E326D_AC9 at any concentration of agonist tested indicating that the efficacy of auto-inhibition by C2b was reduced by the E326D mutation (Fig 2C). The full dataset of this experiment is shown in Fig 3. Closely similar results were obtained with prostaglandin E1 as the agonist (Appendix Fig S1), and further iterations with isoproterenol are provided as Appendix Figs S2 and S3.

The lack of a substantial enhancement of the response to isoproterenol in E326D_AC9C2a is unlikely to be due to the saturation of Glosensor22F as it was apparent with all agonist-induced responses that evoked light-emission well below those elicited by the quality-control forskolin/rolipram stimulus. Comparison of the expression of the AC9 proteins examined here showed that AC9 and AC9C2a were consistently present at higher levels than their E326D counterparts (Fig 4). Thus the cAMP-producing capacities of the E326D mutants are likely underestimated when compared with the wild-type variants. Importantly, the levels of expression of the AC9 proteins lacking C2b were similar to the respective full-length versions (Fig 4). As a quality-control for Glosensor22F expression was run in each well, and as Glosensor 22F is validated for scalar analysis of intracellular cAMP levels (Ayukawa *et al*, 2020; Baldwin *et al*., 2019; Binkowski *et al*, 2011; Felouzis *et al*, 2016; Goulding *et al*, 2018; Hoy *et al*, 2020) it is justified to conclude that the dramatic differences of cellular cAMP levels observed in our experiments largely reflect the respective rates of cAMP production.

**Figure 4.**
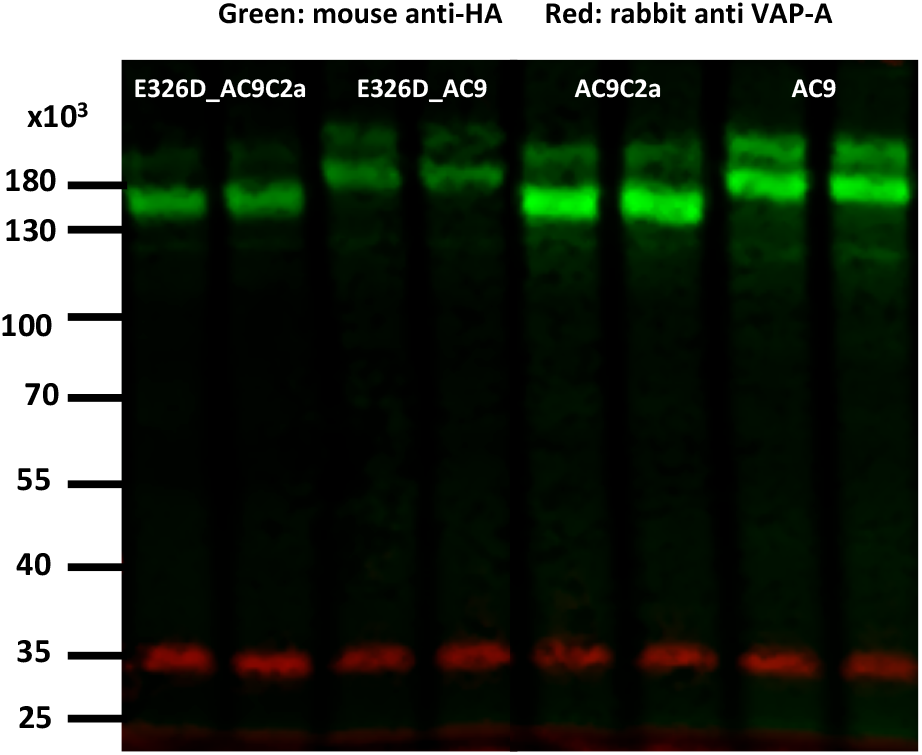
Immunoblot of extracts of crude membranes prepared from HEK293 FT cells transfected with (from left to right, two lanes each): E326D_AC9C2a, E326D_AC9, AC9C2a, AC9. Green bands show reaction with anti_HA, red bands are stained for VAMP-associated protein 33 (VAP-A). The relative intensities of the protein bands reported by Empiria software (LiCor biosciences) were E326D_AC9C2a 0.42, E326D_AC9 0.40, AC9C2a 1.1, AC9 1.0, the blot is representative of three transfections. The numbers on the left indicate the migration of the molecular size markers on the gel.

The data show that in addition to quenching the response to GsR, the C2b domain has an auto-stimulatory effect on AC9. Moreover, perturbation of the coiled-coil of AC9 can differentially modify the influence of the C2b domain, in that it sustains auto-stimulation but appears to supress auto-inhibition. Conformation changes of the coiled-coil could occur under physiological conditions. While fulllength AC9 is resistant to regulation by G-proteins, the trans-membrane arrays may function as cell-surface receptors and/or sensors of the lipid composition of the plasma membrane (Finkbeiner *et al*, 2019; Qi *et al*., 2022; Qi *et al*., 2019; Seth *et al*, 2020). The interfaced helices 6 and 12 of the trans-membrane arrays continue into the cytoplasm as the transducing helices and form the coiled-coil (Qi *et al*., 2019). Overall, this scenario appears similar to the activation of soluble guanylyl cyclase by nitric oxide (Horst *et al*., 2019).

Basal cAMP production by AC9 is inhibited by an intracellular pathway involving Ca^2+^ and calcineurin (Antoni *et al*, 1995; Antoni *et al*, 1998; Cumbay & Watts, 2005; Paterson *et al*, 1995). Hence, it is part of regulated intracellular signalling circuits. In contrast to our results in intact cells, purified preparations of bovine AC9 and AC9C2a (Qi *et al*., 2019) showed no difference in basal enzymatic activity. As the C2b domain may be phosphorylated (ten documented sites) as well as ubiquitinated (3 sites) (Antoni, 2020), it is possible that post-translational modifications are essential for the auto-stimulation observed in HEK293FT cells, and these are likely to have been lost during the multi-step purification process (Qi *et al*., 2019). Indeed, functionally relevant activation of AC9 by protein kinase cascades independently of Gs has been reported in neutrophil granulocytes (Liu *et al*, 2010). A further possibility is that auto-stimulation may require additional protein(s) lost during purification.

The autoinhibitory motif of C2b that occludes AC9 when in complex with Gsα is well delineated (Pálvölgyi *et al*., 2018; Qi *et al*., 2019). With respect to how C2b might stimulate AC9 activity, indirect evidence points to the forskolin binding-pocket (Qi *et al*., 2022; Tang & Hurley, 1998). First, the cryo-electronmicroscopic map of bovine AC9 shows that the C2b domain is capable of short-distance interactions with residues in the forskolin binding-pocket (Qi *et al*., 2019). Second, database analysis of the phylogenetic development of AC9 reveals that a long (>100 amino acid residues) C2b domain containing the highly conserved auto-inhibitory motif (Pálvölgyi *et al*., 2018) only features in vertebrate AC9-s. Simultaneously, a well-defined, “low-reactivity to forskolin” configuration (Tang & Hurley, 1998; Yan *et al*, 1998) of the C2a catalytic domain also emerges. These features are already present in lamprey as well as hagfish AC9, the two earliest vertebrate species alive today. In contrast, the invertebrate homologs of AC9, including those of the chordate amphioxiform species, have short C2b domains and their C2a domains are in the “high-reactivity to forskolin” configuration. Hence, it seems reasonable to suggest that the “low-reactivity to forskolin” configuration of the C2a domain of AC9 is instrumental to autostimulation by the C2b domain.

### Final summary

AC9 features auto-stimulation of basal activity as well as auto-inhibition of Gsα activation by its isoform-specific C2b domain. These facets of auto-regulation may be differentially modulated through the coiled-coil of the signalling helices. Last but not least, AC9 may be recruited into the GsR regulatory circuit by proteolytic cleavage of the COOH-terminal C2b domain, which markedly lowers basal activity and enables the response to Gsα (Pálvölgyi *et al*., 2018). Figure 5 shows the three modes of operation of AC9 discussed above. In principle, all three may occur simultaneously in the same cell. As AC9 is widely distributed in the body, these features of intracellular cAMP signalling are bound to be relevant in several organ systems.

**Fig 5:**
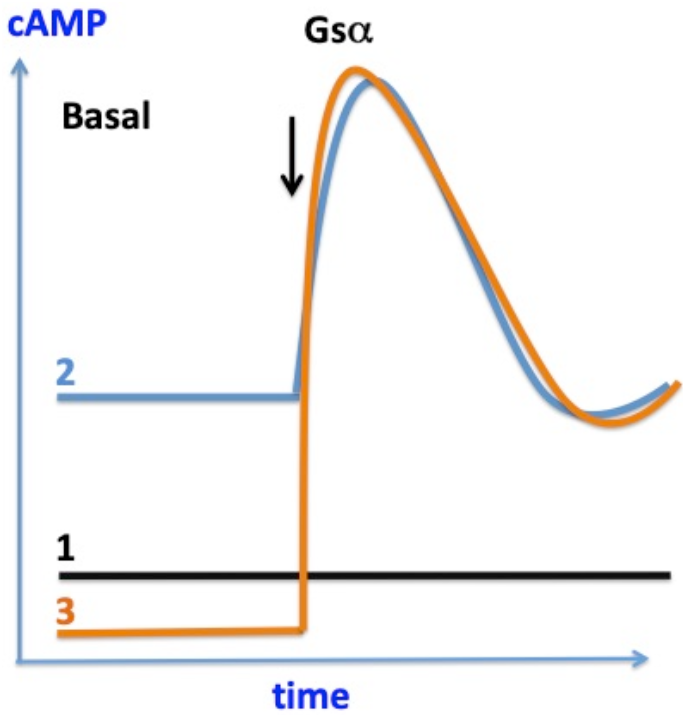
Three modes of operation of AC9 as indicated by this study and previous work. Mode 1: High basal activity supported by the C2b domain and quenching of activation “occluded state” in the presence of activated Gsα, as shown in previous work. Importantly, the production of cAMP by AC9 is inhibited by intracellular free Ca^2+^ through calcineurin. Mode 2: Conformation change at the coiled coil (E326D mutation in this study) enhances basal activity dependent on C2b and relieves the quenching of the activation by Gsα. Thus the enzyme may summate input to the coiled-coil and the stimulation by Gsα Mode 3: Removal of the C2b domain markedly reduces basal activity and enables stimulation by Gsα, i.e. AC9 now resembles a conventional adenylyl cyclase stimulated by Gsα.

## Materials and Methods

### cDNA constructs

Human AC9 tagged N-terminally with haemagglutinin antigen (HA) and C-terminally with FLAG in pcDNA3.1 (AC9) was a gift of Dr. Adrienn Pálvölgyi (Egis PLC, Budapest, Hungary). This construct was used for site-directed mutagenesis with Phusion polymerase (NEB, Ipswich, MA.,U.S.A.) following previously published protocols (Xia *et al*, 2015). Two mutations were produced - L323S and E326D. Both amino acid residues are part of the interface of the coiled-coil of AC9 (Fig 2B and C). Subsequently, the C2b domain was removed from AC9 and E326D_AC9 (Pálvölgyi *et al*., 2018) to encode AC9_C2a and E326D_AC9C2a, respectively. The sites of the mutations as well as the N- and C-terminal coding sequences were verified by Big Dye sequencing.

### Cells

Fast-growing human embryonic kidney 293T cells (HEK293FT) were maintained in DMEM 10% FBS (v/v) and passaged at 5-7 day intervals with TrypleExpress (Invitrogen GIBCO, Paisley, UK,) to detach the cells. About 2 million cells in 1.6 ml of growth medium were mixed with 4 μg of AC9 cDNA or 2μg of pcDNA3 plus 2 μg of GlosensorF22 (Binkowski *et al*., 2011) pre-complexed with Lipofectamine 2000 in 400 μl OPTIMEM. GlosensorF22 encodes a firefly luciferase-based biosensor that, when provided with its substrate luciferin, emits light in proportion to the amount of cAMP bound to it (Binkowski *et al*., 2011).

### Measurement of cellular cAMP levels

After 48h, the transfected cells were plated in poly-l-lysine coated, 96-well white tissue culture plates (Greiner, Stonehouse, UK) at 10^5^ cells per well and incubated as above for 24 h. Subsequently the cells were depleted of serum in DMEM for 60 min and incubated in Hank’s balanced salt solution containing 1 mM MgSO4, 1.5 mM CaCl_2_, 10 mM HEPES pH7.4 and 1 mM beetle luciferin (Promega, Southampton, UK) at 32°C for a further 60 min. The plates were transferred to a BMG Lumistar Omega plate-reader and the luminescence signal was recorded at 32°C from each well at 2 min intervals. Usually a further 30 min were required for the basal light signal to stabilize. Drug treatments were added from a 12-channel handheld pipette. Due to the inherent variability of transient transfections each well received a mixture of 5 μM forskolin (LC labs, Woburn, MA, U.S.A.) and 100 μM rolipram (Insight Biotech, Wembley, U.K.) at the end of the recording as a quality control stimulus. This standardization was possible because AC9 is not stimulated by 5 μM forskolin even when stimulated by Gsα (Baldwin *et al*., 2019; Qi *et al*., 2022; Qi *et al*., 2019). As the glosensor response at high levels of cAMP becomes non-linear and eventually saturates (Binkowski *et al*., 2011), the concentration-response curves may be flattened at high concentrations of isoproterenol. This is likely to be the case when the isoproterenol response (largely generated by the transfected AC9 variant) is higher than the standardizing stimulus that is largely produced by the host cell adenylyl cyclases.

### Immunoblots

Expression of HAFLAG-tagged AC9 proteins was examined by SDS-PAGE and immunoblotting of extracts prepared from crude membranes of transfected HEK293 FT cells as reported previously (Antoni *et al*., 1998; Pálvölgyi *et al*., 2018). Protein blots were reacted with the 12CA5 Anti-HA (Abcam, Cambridge, UK) or M2 anti-Flag mouse monoclonal antibodies (Sigma-Aldrich, Poole, Dorset UK) in conjunction with rabbit anti-VAP-A (gift of Dr Paul Skehel) (Skehel *et al*, 2000) as a sample loading control marker. Secondary IRDye 680RD-tagged anti-mouse and IRDye 800CW-tagged anti-rabbit goat IgGs were from LiCor Biosciences (Lincoln, NE, USA) with fluorescence read-out in a LiCor Odyssey imager. Only HA-tag staining was used for quantification as the staining with Anti-FLAG M2 antibody appeared to show context dependence.

### Database searches

The NCBI Genbank and the Wellcome Trust Ensemble Genome Browser servers were used to find AC9 related sequences by BLAST searches. Protein sequence alignments were carried out with the Clustal Omega web-application on the European Bioinformatics Institute server (Madeira *et al*, 2022). The cryo-electronmicroscopic maps of bovine AC9 were downloaded from the RSCB Protein Data Bank server.

## Acknowledgments

This study was funded by the University of Edinburgh. We would like thank Miss Heather McClafferty for help with cell lines and various reagents, Dr Paul LeTissier for access to a luminometer, Prof Mike Shipston, Dr Paul Skehel, Dr Sutherland MacIver and Dr Tamás Balla for helpful discussions, Dr Mandy Jackson and Dr Paul Skehel for access to laboratory facilities and reagents.

## Appendix PDF: Expanded view 3 figures

Results obtained with three different batches of transfected HEK293FT cells are shown below.

**Fig. S1.**
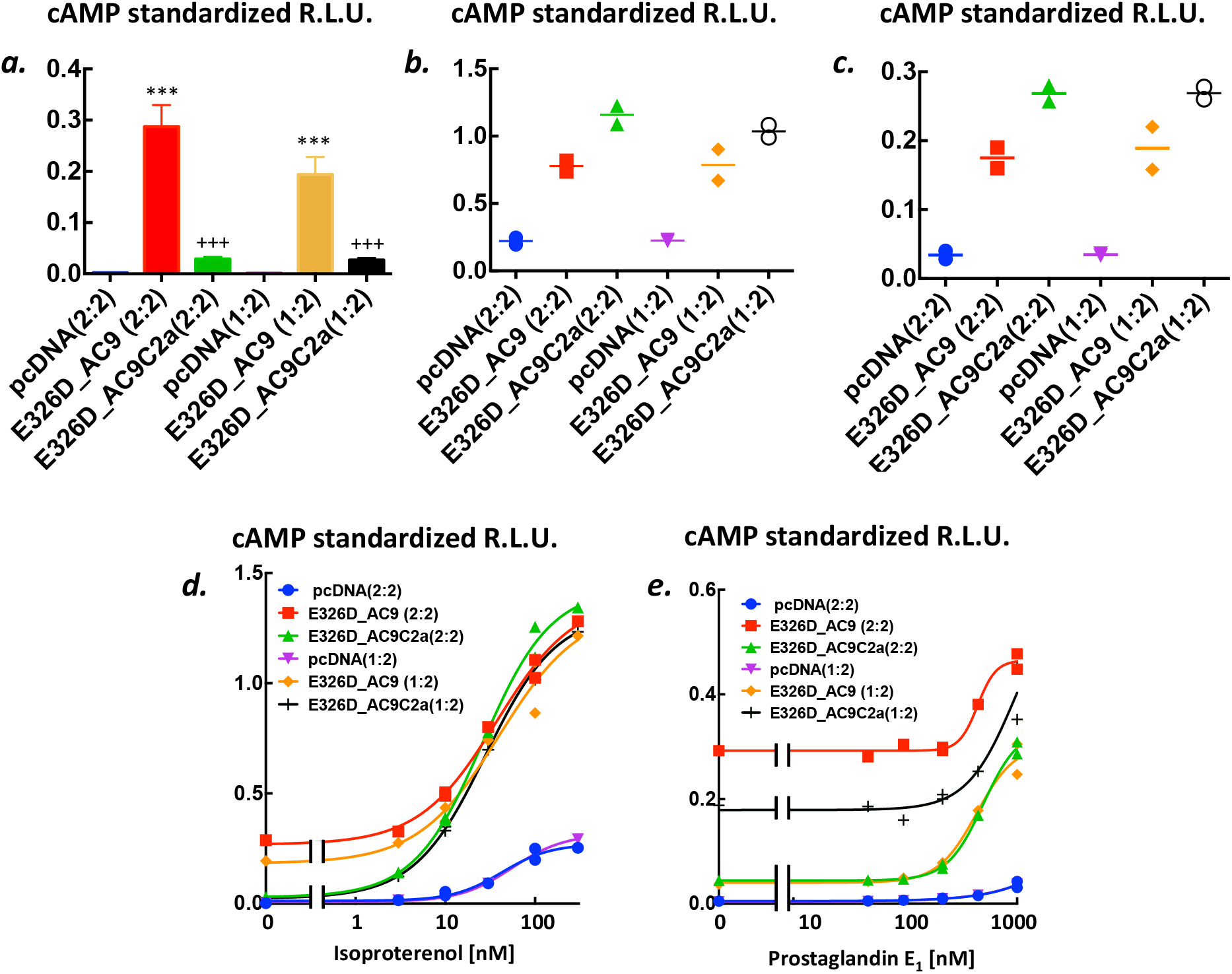
Mutation E326D of the coiled-coil markedly increases basal and agonist-induced cAMP levels. The increase of basal levels is dependent on the presence of the C2b domain. a) Basal intracellular levels of cAMP reported by glosensor22F bioluminescence in HEK293FT cells transiently transfected with 2μg glosensor22F plasmid and 1 (1:2) or 2 (2:2) μg of the plasmids of interest that encoded pcDNA3, E326D_AC9 or E326D_AC9C2a, respectively. Data are the average of the apparent plateau reached at 25-30 min of incubation (see Fig 3 of main paper) standardized in each well by the peak of the response elicited by 5μM forskolin and 100μM rolipram administered at the end of the experiment. Data are means ±S.D. n = 14/group. As the variances of the six groups were significantly different, the statistical analysis was carried out after log-transformation of the data. One-way ANOVA F(5,78)=1554 P<0.0001, followed by Tukey’s *post-hoc* multiple comparison test: + + + P<0.0001 significantly different from the respective pcDNA3 group, *** P<0.0001 when compared with the respective E326D_AC9C2A group. b) The increment over the respective basal levels induced by 100 nM isoproterenol calculated by averaging three consecutive time-points once the peak level of bioluminescence was reached (see Fig 3 of main paper). The relative light-units were standardized in each well by the peak of the response elicited by 5μM forskolin and 100μM rolipram administered at the end of the experiment. All points are shown, the lines represent the means. c) The increment over the respective basal levels induced by 1000 nM prostaglandin E1. Other details as in Suppl fig 1b. d) Concentration-response to isoproterenol. All measured points are shown. Nonlinear four-parameter regression was used to fit the curves in Graphpad Prism v.6. The calculated EC50s (range 28-55 nM) and the maximal responses were not different at alpha = 0.05. e) Concentration-response to prostaglandin E_1_. All measured points are shown. As the maximum of the curve could not be estimated, EC50 values were not calculated.

**Fig. S2.**
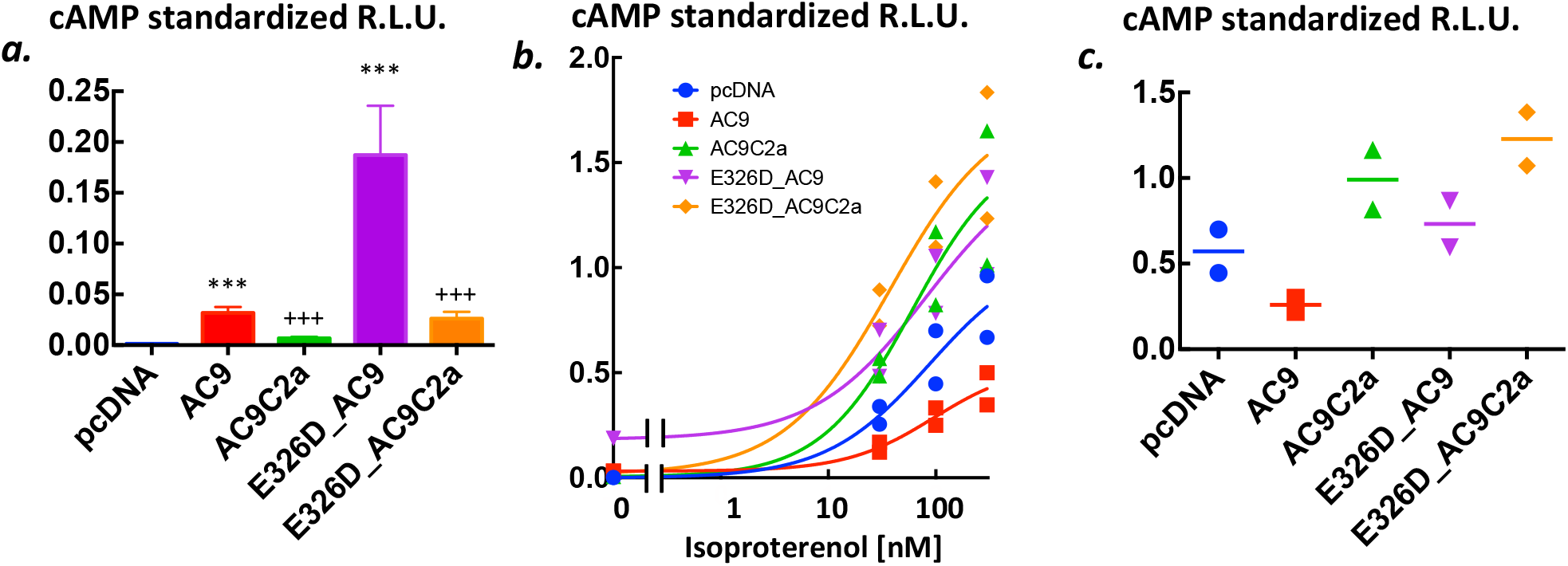
Auto-stimulation of basal and auto-inhibition of isoproterenol-evoked cAMP production by the C2b domain of AC9. a) Basal intracellular cAMP levels reported by glosensorF22 bioluminescence in HEK293FT cells transiently expressing pcDNA3, AC9, AC9C2a, E326D_AC9 or, E326D_AC9C2a. Data are the average of the apparent plateau reached at 25-30 min of incubation (see Fig 3 of main paper) standardized in each well by the peak of the response elicited by 5μM forskolin and 100μM rolipram administered at the end of the experiment. Data are means ± S.D. n = 6/group. As the variances of the five groups were significantly different, statistical analysis was carried out after log-transformation of the data. One-way ANOVA, F(4,25) = 320 P<0.0001. Tukey’s *post-hoc* multiple comparison test: *** P<0.001 *vs*. respective variant lacking the C2b domain. + + + P<0.0001 *vs*. pcDNA3 group. b) Concentration-response to isoproterenol from the cells described in a). All measurements of the standardized peak responses are shown. Curves were fitted by non-linear four-parameter regression with variable slope in Graphpad Prism v.6. As the maximum of the curves could not be estimated reliably, no EC50 values are reported. c) The increment over the respective basal levels induced by 100 nM isoproterenol calculated by averaging three consecutive time-points once the peak level of bioluminescence was reached (see Fig 3 of main paper). The relative light-units were standardized in each well by the peak of the response elicited by 5μM forskolin and 100μM rolipram administered at the end of the experiment. All points are shown, the lines represent the means.

**Fig. S3.**
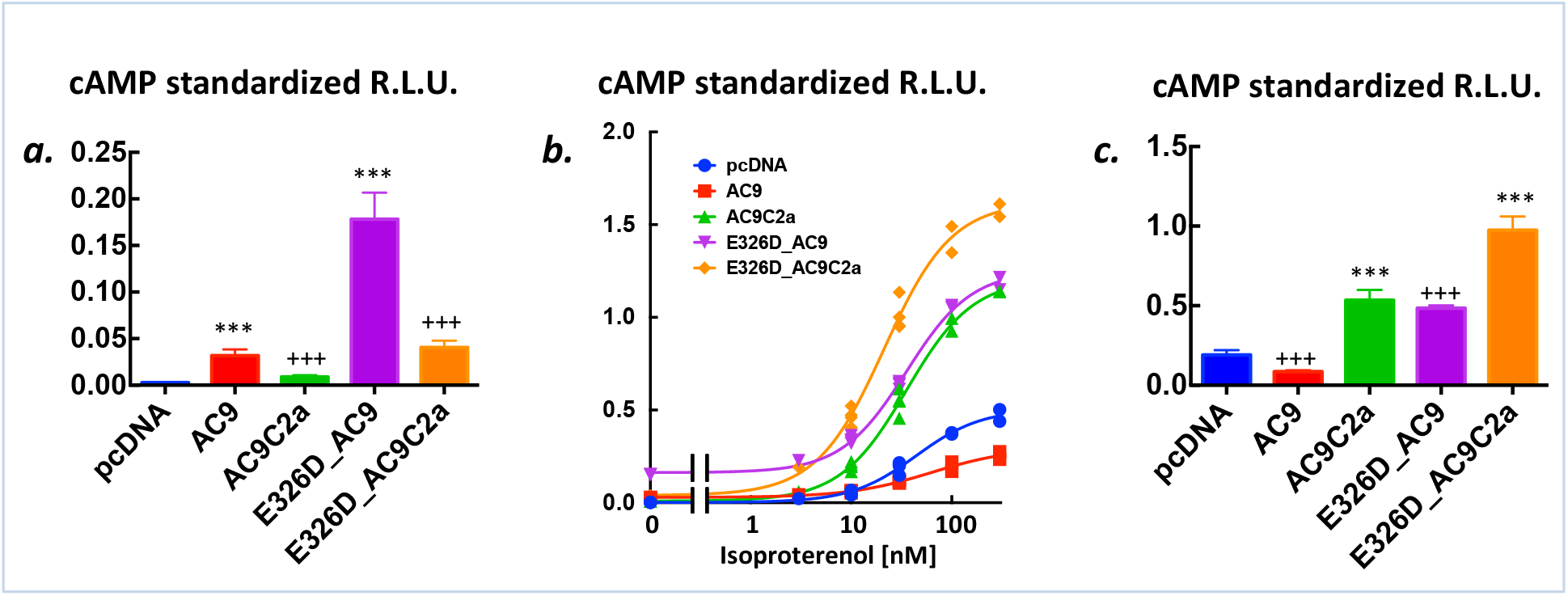
Auto-stimulation of basal and auto-inhibition of isoproterenol-evoked cAMP production by the C2b domain of AC9. a) Basal cAMP levels, all as in Suppl Fig 2a) except: Data are means ± S.D. n = 14/group. As the variances of the five groups were significantly different, statistical analysis was carried out after log-transformation of the data. One-way ANOVA, F(4,65) = 935 P<0.0001. Tukey’s *post-hoc* multiple comparison test: *** P<0.001 *vs*. respective variant lacking the C2b domain. + + + P<0.0001 *vs*. pcDNA3 group. b) Concentration response to isoproterenol from the cells described in a). All measurements of the standardized peak responses are shown. Curves were fitted by non-linear four-parameter regression with variable slope in Graphpad Prism v.6. The EC50 of the E326D_AC9C2a (21 nM) was significantly lower (95% confidence intervals) than that of the other four groups – range 35-65 nM. The maximal responses were E326D_AC9C2a> E326D_AC9= AC9C2a>pcDNA3>AC9 where > denotes statistical significance at P<0.05, as indicated by the 95% confidence intervals calculated by the non-linear regression algorithm. c) The increment of cAMP levels above the respective basal levels at 30 nM isoproterenol. Means ± S.D. n=4/group. Data reduction protocol as described in Suppl Fig 1b. One-way ANOVA F(4,15) = 186.1 P<0.0001. Tukey’s *post-hoc* multiple comparison test: *** P<0.001 *vs*. respective full-length variant. +++ P<0.0001 *vs*. pcDNA3 group. The results for 10 nM isoproterenol were closely similar.

